# Neuromuscular control of gait stability in older adults is adapted to environmental demands but not improved after standing balance training

**DOI:** 10.1101/2020.10.28.358788

**Authors:** Leila Alizadehsaravi, Sjoerd M. Bruijn, Wouter Muijres, Ruud A.J. Koster, Jaap H. van Dieën

## Abstract

Balance training aims to improve balance and transfer acquired skills to real-life tasks and conditions. How older adults adapt gait control to different conditions, and whether these adaptations are altered by balance training remains unclear. We investigated adaptations in neuromuscular control of gait in twenty-two older adults (72.6 ± 4.2 years) between normal (NW) and narrow-base walking (NBW), and the effects of a standing balance training program shown to enhance unipedal balance control in the same participants. At baseline, after one session and after 3-weeks of training, kinematics and EMG of NW and NBW on a treadmill were measured. Gait parameters and temporal activation profiles of five synergies extracted from 11 muscles were compared between time-points and gait conditions. No effects of balance training or interactions between training and walking condition on gait parameters or synergies were found. Trunk center of mass (CoM) displacement and velocity (vCoM), and the local divergence exponent (LDE), were lower in NBW compared to NW. For synergies associated with stance of the non-dominant leg and weight acceptance of the dominant leg, full width at half maximum (FWHM) of the activation profiles was smaller in NBW compared to NW. For the synergy associated with non-dominant heel strike, FWHM was greater in NBW compared to NW. The Center of Activation (CoA) of the activation profile associated with dominant leg stance occurred earlier in NBW compared to NW. CoAs of activation profile associated with non-dominant stance and non-dominant and dominant heel strikes were delayed in NBW compared to NW. The adaptations of synergies to NBW can be interpreted as related to a more cautious weight transfer to the new stance leg and enhanced control over CoM movement in the stance phase. However, control of mediolateral gait stability and these adaptations were not affected by balance training.

## Introduction

Falls in older adults mostly occur during walking [1]. Therefore, skills acquired during standing balance training should transfer to gait and improve gait stability [2]. While on one hand effects of balance training have been described as task specific [3], on the other hand, transfer from standing balance training to gait stability has been suggested by improved clinical balance scores and gait parameters [4,5]. Consequently, the existence of skill transfer from standing balance training as well as the mechanisms underlying such transfer, if present, are insufficiently clear.

Increased variability and decreased local dynamic stability of steady-state gait were shown to be associated with a history of falls in older adults [6]. From a mechanical perspective, larger mediolateral center of mass excursions and velocities would be expected to cause an increased fall risk [7] and both these parameters as well as their variability are larger in older than young adults [8]. When facing environmental challenges, such as when forced to walk with a narrow step width, individuals need to adapt their gait. Older adults show more pronounced adaptations to narrow-base walking compared to young adults [8], possibly because they are more cautious in the presence of postural threats [9]. Transfer of standing balance training to gait would be expected to result in increased gait stability, decreased CoM displacement and velocity, and decreased CoM displacement variability. In addition, an interaction between training and stabilizing demands may be expected. Increased confidence after training may result in less adaptation to a challenging condition. On the other hand, balance training may enhance the ability to adapt to challenging conditions.

The central nervous system is thought to simplify movement by activating muscles in groups, called muscle synergies, with the combination of synergies shaping the overall motor output [10,11]. Muscle synergies consist of time-dependent patterns (activation profiles) and time-independent factors (muscle weightings). Human gait has been described with four to eight muscle synergies [12–14] and reactive balance control was found to have four shared synergies with walking [14], which could be important for transfer from balance training to gait. Due to aging and changes in sensory and motor organs, adapted synergies are likely required to maintain motor performance [15,16]. Synergy analyses of gait revealed either fewer synergies in older adults than in young adults [17] or no differences [18]. Motor adaptation is assumed to result from altering synergies in response to task and environmental demands [19,20]. For example, widened activation profiles appear to be used to increase the robustness of gait in the presence of unstable conditions or unpredictable perturbations [20,21]. Long-term balance training might alter synergies in gait, and adaptation of synergies to task demands as has been shown in dancers [22,23] to achieve the alterations in CoM kinematics.

We investigated the adaptations in neuromuscular control of gait in older adults between normal and narrow-base walking, and the effect of short- and long-term standing balance training on this. To this aim, we used data from a previous study on standing balance training, from which we previously reported positive effects of training on standing balance robustness and performance, both after a single training session and after three weeks of training [24]. Here, we evaluate skill transfer to normal walking and narrow-base walking on a virtual beam, both on a treadmill. We used foot placement error to assess performance of narrow-base walking [25]. We focused on mediolateral balance control, as larger mediolateral instability has been shown to be associated with falls in older adults [26,27] and beam walking challenges mediolateral stability. We calculated the CoM displacement and CoM displacement variability, CoM velocity and the LDE as measures of gait stability and extracted muscle synergies to characterize effects on the neuromuscular control of gait and of adaptations to narrow-base walking.

## Methods

The methods described here in part overlap with our previous paper [24], as data were obtained in the same cohort.

### Participants

Twenty-two older (72.6 ± 4.2 years old; mean ± SD, 11 females) healthy volunteers participated in this study. Participants were recruited through a radio announcement, contacting older adults who previously participated in our research, flyers and information meetings. Individuals with obesity (BMI > 30), cognitive impairment (MMSE<24), peripheral neuropathy, a history of neurological or orthopedic impairment, use of medication that may negatively affect balance, inability to walk for 4 minutes without aid, and performing sports with balance training as an explicit component (e.g., Yoga or Pilates) were excluded. All participants provided written informed consent before participation and the procedures were approved by the ethical review board of the Faculty of Behavioural & Movement Sciences, VU Amsterdam (VCWE-2018-171).

### Experimental procedures

Participants completed an initial measurement to determine baseline values (Pre), a single-session balance training (30-minutes), a second measurement (Post1) to compare to baseline to assess short-term training effects, a 3-week balance training program (9 sessions x 45 minutes training), and a third measurement (Post2) to compare to baseline to assess of long-term training effects (Fig 1).

**Fig 1.**
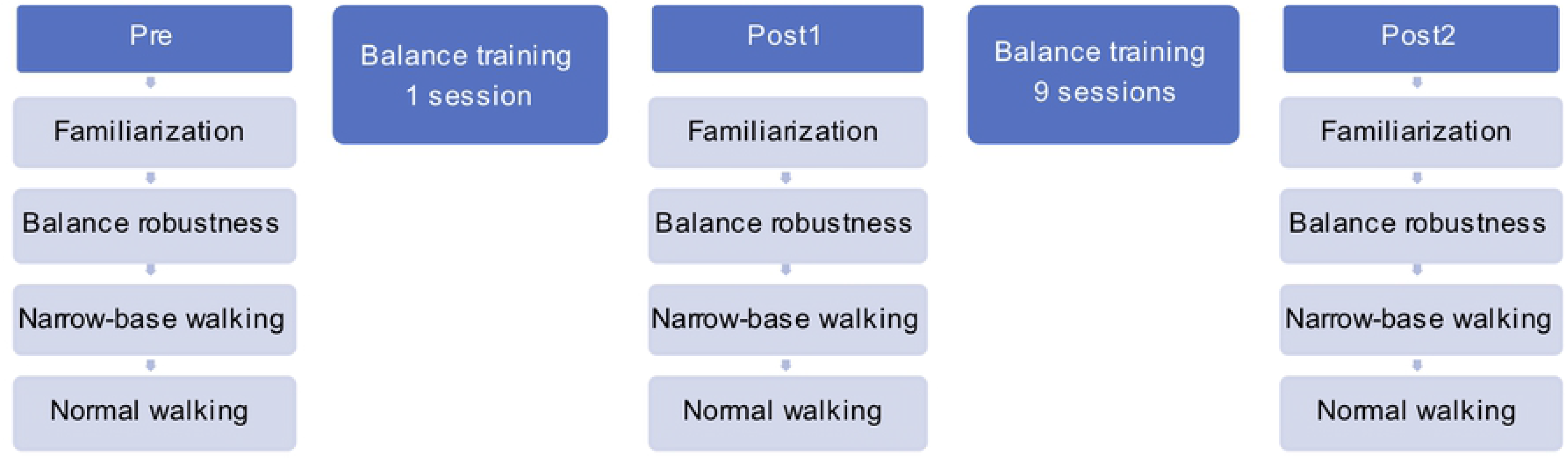
Block diagram of the study; training and gait assessment.

The measurements consisted of one experimental condition on a robot-controlled platform (balance robustness) and two experimental conditions performed on a treadmill: virtual-narrow-base walking (Fig 2) and normal walking.

**Fig 2.**
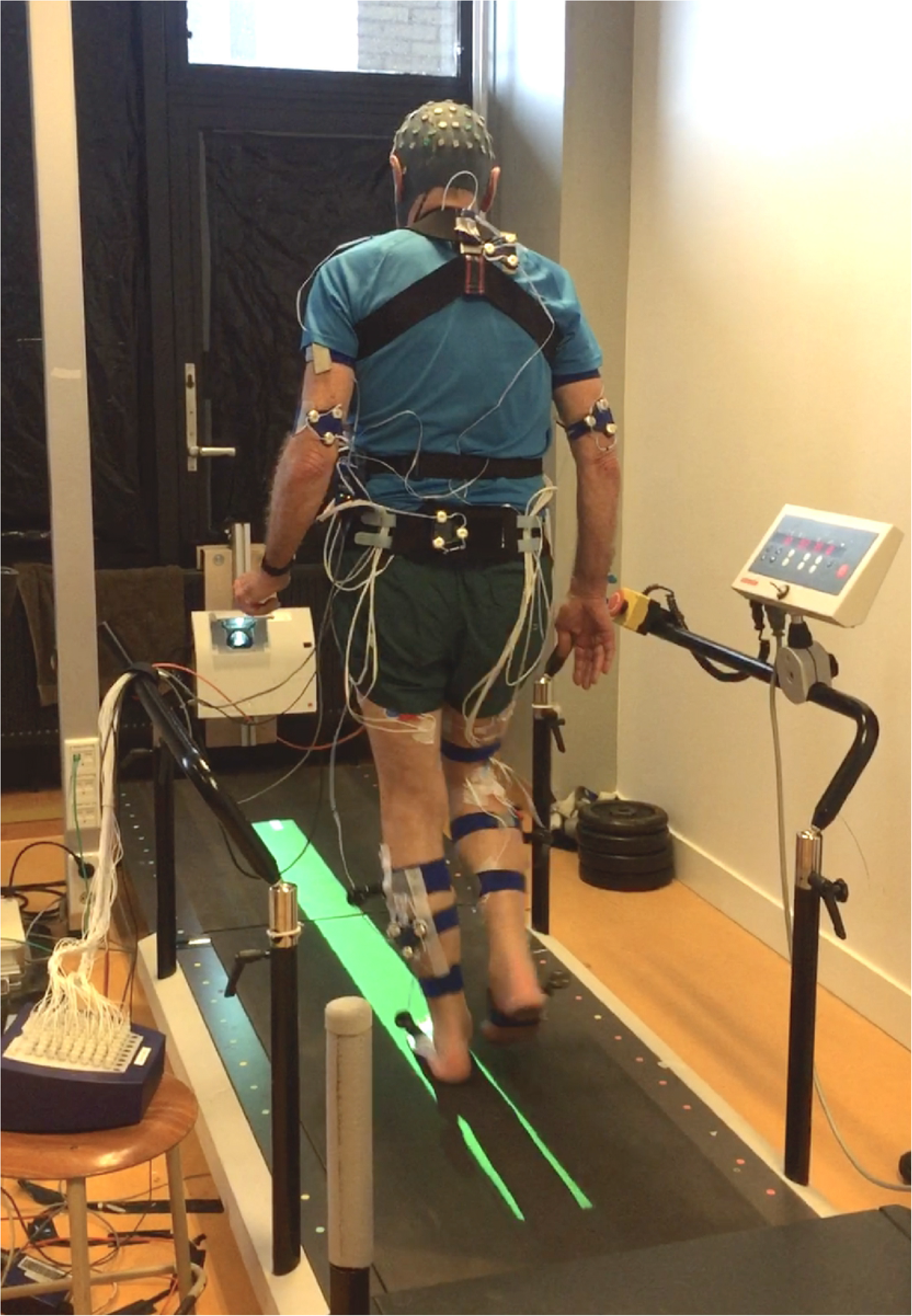
Narrow-base walking on a treadmill.

The training sessions consisted of exercises solely focused on unipedal balancing with blocks of 40-60 second exercises in which balance was challenged by different surface conditions, static vs dynamic conditions, perturbations, and dual tasking (e.g. catching, throwing and passing a ball) [28]. Participants performed the exercises in a group of two (except for the first, individual session) and always under supervision of the physiotherapist in our research team.

### Instrumentation and data acquisition

Balance robustness and performance were evaluated using a custom-made balance platform controlled by a robot arm (HapticMaster, Motek, Amsterdam, the Netherlands) and results were reported previously [24]. To quantify transfer to gait, participants were instructed to walk for 4.5 minutes at a constant speed of 3.5 km/h on a treadmill with an embedded force plate. For safety reasons, handrails were installed on the either side of the treadmill, and an emergency stop button was placed within easy reach (MotekForcelink, Amsterdam, the Netherlands). We assessed walking in two conditions, normal walking and narrow-base walking, in a randomized order, with a minimum of 2 minutes seated rest in between conditions. In narrow-base walking, participants were instructed to placing their entire foot inside the beam as accurately as possible over a green light-beam path (12 cm width) projected in the middle of the treadmill (Bonte Technology/ForceLink, Culemborg, The Netherlands) [25].

Kinematics data were obtained by two Optotrak 3020 camera arrays at 50 Hz (Northern Digital, Waterloo, Canada). 10 active marker clusters (3 markers each) were placed on the posterior surface of the thorax (1), pelvis (1), arms (2), calves (4), and feet (2) (Fig 2). Positions of anatomical landmarks were digitized by a 4-marker probe and a full-body 3D-kinematics model of the participant was formed relating clusters to the neighboring landmarks [29]. The position of the foot segments was obtained through cluster markers on both feet, digitizing the medial and lateral aspects of the calcaneus, and the heads of metatarsals one and five [25]. Additionally, to calculate the foot placement error in narrow-base walking, position and orientation of the projected beam was determined by digitizing the four outer bounds of the beam on the treadmill.

Surface electromyography (EMG) data were recorded from 11 muscles; 5 unilateral muscles of the dominant leg: tibialis anterior (TAD), vastus lateralis (VLD), lateral gastrocnemius (GLD), soleus (SOD), peroneus longus (PLD) and, 6 bilateral muscles: rectus femoris (RFD, RFN), biceps femoris (BFD, BFN) and gluteus medius (GMD, GMN) muscles. Bipolar electrodes were placed in accordance with SENIAM recommendations [30]. EMG data were sampled at a rate of 2000 Hz and amplified using a 16-channel TMSi Porti system (TMSi, Twente, The Netherlands). The dominant leg was the leg preferred for single-leg stance. Focus was on this leg, because we extensively assessed unipedal balance control on this leg as reported earlier [24].

### Data analysis

#### Gait events

The first 30 seconds of all gait trials were removed, to discard the habituation phase. Heel-strikes were detected through a peak detection algorithm based on the center of pressure [31]. This algorithm proved to be precise when the center of pressure moved in a butterfly pattern. However, for narrow-base walking, the feet share a common area in the middle of the treadmill, therefore, identification of which leg touched the surface was problematic. Hence, heel-strikes were detected based on the center of pressure peak detection, but the associated leg was identified based on kinematic data of the foot marker. 160 strides per participant per condition were used to calculate all gait variables (i.e. stability variables and muscle synergies).

#### Gait stability

To evaluate gait performance, foot placement errors were determined as the mean mediolateral distance of the furthest edge of the foot from the edge of the beam. If the foot was within the beam the error equals zero.

The trajectory of the center of mass (CoM) of the trunk was estimated from mediolateral trunk movement [32,33]. As gait stability variables, we calculated mean and standard deviation of the peak-to-peak mediolateral trunk CoM displacement and mean of CoM velocity per stride. In addition, local dynamic stability was evaluated using the local divergence exponent, LDE, based on Rosenstein’s algorithm [34,35]. We used the time normalized time-series (i.e. 160 strides of data were time normalized to 16000 samples, preserving between stride variability) of trunk vCoM to reconstruct a state space with 5 embedding dimensions at 10 samples time delay [33]. The divergence for each point and its nearest neighbor was calculated and the LDE was determined by a linear fit over half a stride to the averaged log transformed divergence.

#### Muscle synergies

EMG data were high-pass (50 Hz, bidirectional, 4th order Butterworth) [20] and notch filtered (50 Hz and its harmonics up to the Nyquist frequency, 1 Hz bandwidth, bidirectional, 1st order Butterworth). The filtered data were Hilbert transformed, rectified and low-pass filtered (10 Hz, bidirectional, 2nd order Butterworth). Each channel was normalized to the maximum activation obtained for an individual per measurement point per trial. Synergies were extracted from 11 muscles using non-negative matrix factorization. Five synergies were extracted from the whole dataset, to account for a minimum of 85% of the variance in the EMG data (Fig 3). It has been shown that perturbations during walking change the temporal activation profiles as compared to normal walking, while muscle weightings are preserved [36]. Therefore, in the current study we fixed muscle weightings between conditions and time-points. These muscle weightings were extracted from the concatenated EMG data of both conditions at all time-points. This allowed for objective comparison of synergy activation profiles between normal and narrow-base walking and between time-points. Consequently, the time-normalized EMG data of the muscles **W_11 x (2 x 100 x 160),_** was factorized to two matrices: time-invariant muscle weightings, **H_11 x 5_**, and temporal activation profiles of the factorization, **M_5 x (2 x 100 x 160)_**, where 11 was the number of muscles, 5 the number of synergies, 2 the number of conditions, 100 the number of samples in each stride and 160 the number of strides. Afterwards, we reconstructed the temporal activation profiles using pseudo-inverse multiplication, for the comparison of activation profiles between conditions and time-points.

**Fig 3.**
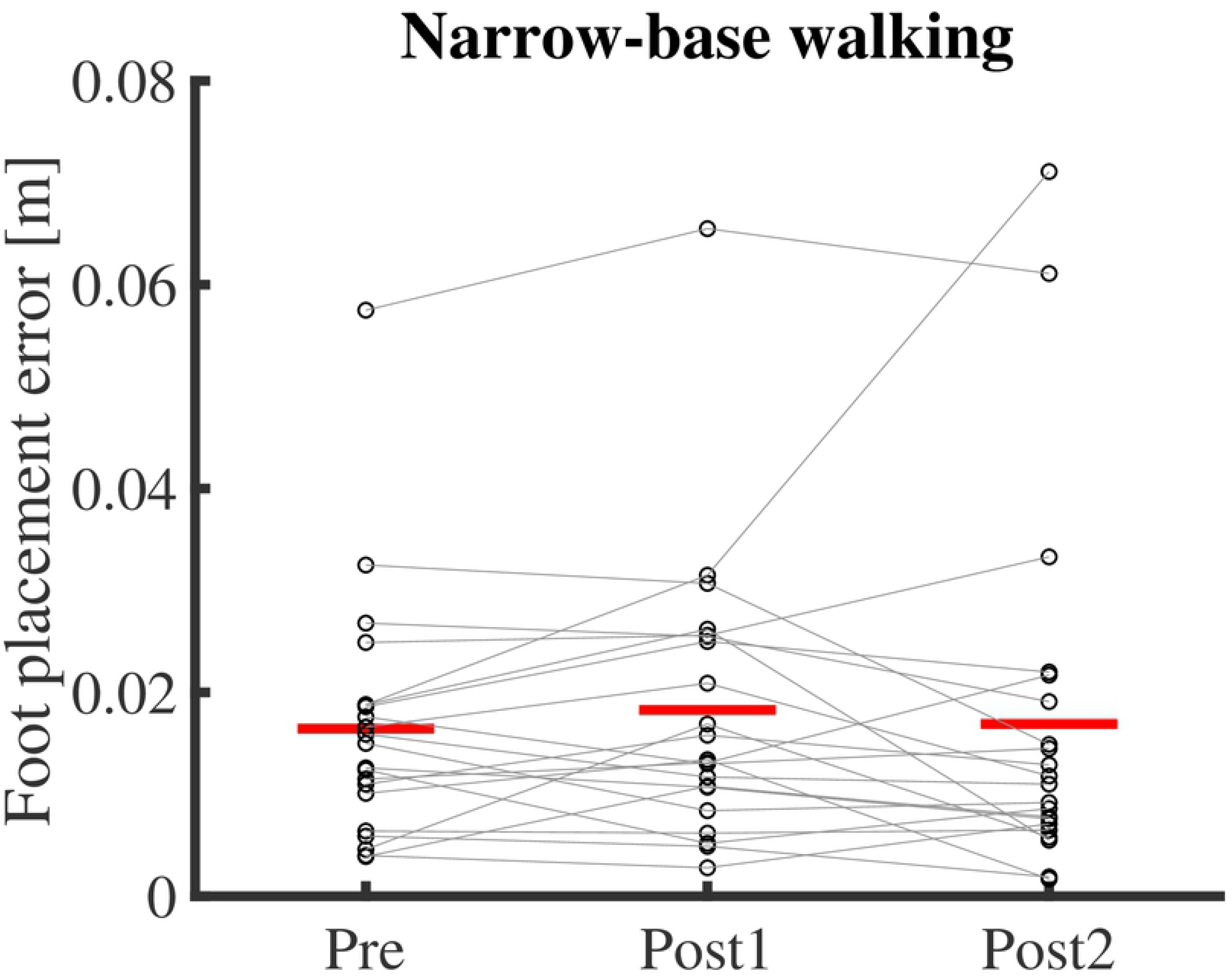
Foot placement error in narrow-base walking at time-points Pre, Post1 and Post2. Thin lines represent individual subject data. Red horizontal lines indicate means over subjects.

To compare activation profiles, we evaluated the full width at half maximum, FWHM, per stride for each activation profile (defined as the number of data points above the half maximum of activation profile, after subtracting the minimum activation [37]). In addition, we evaluated the center of activity, CoA, per stride defined as the angle of the vector that points to the center of mass in the activation profile transformed to polar coordinates [20,38]. FWHM and CoA were averaged over 160 strides per participant per condition. For CoA data, circular averaging was used.

### Statistics

Effects of time-point (Pre, Post1, Post2) on foot placement errors were tested using a one-way repeated measures ANOVA. Post-hoc comparisons (paired sample t-tests), with Holm’s correction for multiple comparisons were performed to investigate the effect of short- and long-term training (Pre vs Post1 and Pre vs Post2, respectively).

Two-way repeated-measures ANOVAs were used to identify main effects of time-point (Pre, Post1, Post2) and condition (normal and narrow-base walking) on trunk kinematics CoM displacement, CoM displacement variability, vCoM and LDE, as well as, on the FWHM. When the assumption of sphericity was violated, the Greenhouse-Geisser method was used. In case of a significant effect of time-point, or an interaction of time-point x condition, post hoc tests with Holm’s correction for multiple comparisons were performed. To identify effects on CoA, parametric two-way ANOVA for circular data was used using the Circular Statistic MATLAB toolbox [39]. In all statistical analyses α = 0.05 was used.

## Results

One participant was not able to perform the treadmill walking trials for the full duration and data for this participant were excluded.

### Gait performance

In contrast with robustness and performance in unipedal balancing [24], performance in narrow-base walking, as reflected in foot placement errors, did not did not improve as a result of training (F_1.267,25.347=_ 0.31, p = 0.63; Fig 3).

Training did also not significantly affect CoM displacement, CoM displacement variability, and vCoM (F_2,40=_ 2.729, p = 0.082; F_2,40=_ 0.469, p = 0.628; F_2,40=_ 2.024, p = 0.145). Condition significantly affected all three variables, with lower CoM and vCoM (F_1,20=_ 96.007, p < 0.001; F_1,20=_ 168.26, p < 0.001, respectively, Fig 4), but larger CoM variability (F_1,20=_ 4.678, p = 0.042), in narrow-base compared to normal walking. No significant interactions of time-point x condition were found (p > 0.05).

**Fig 4.**
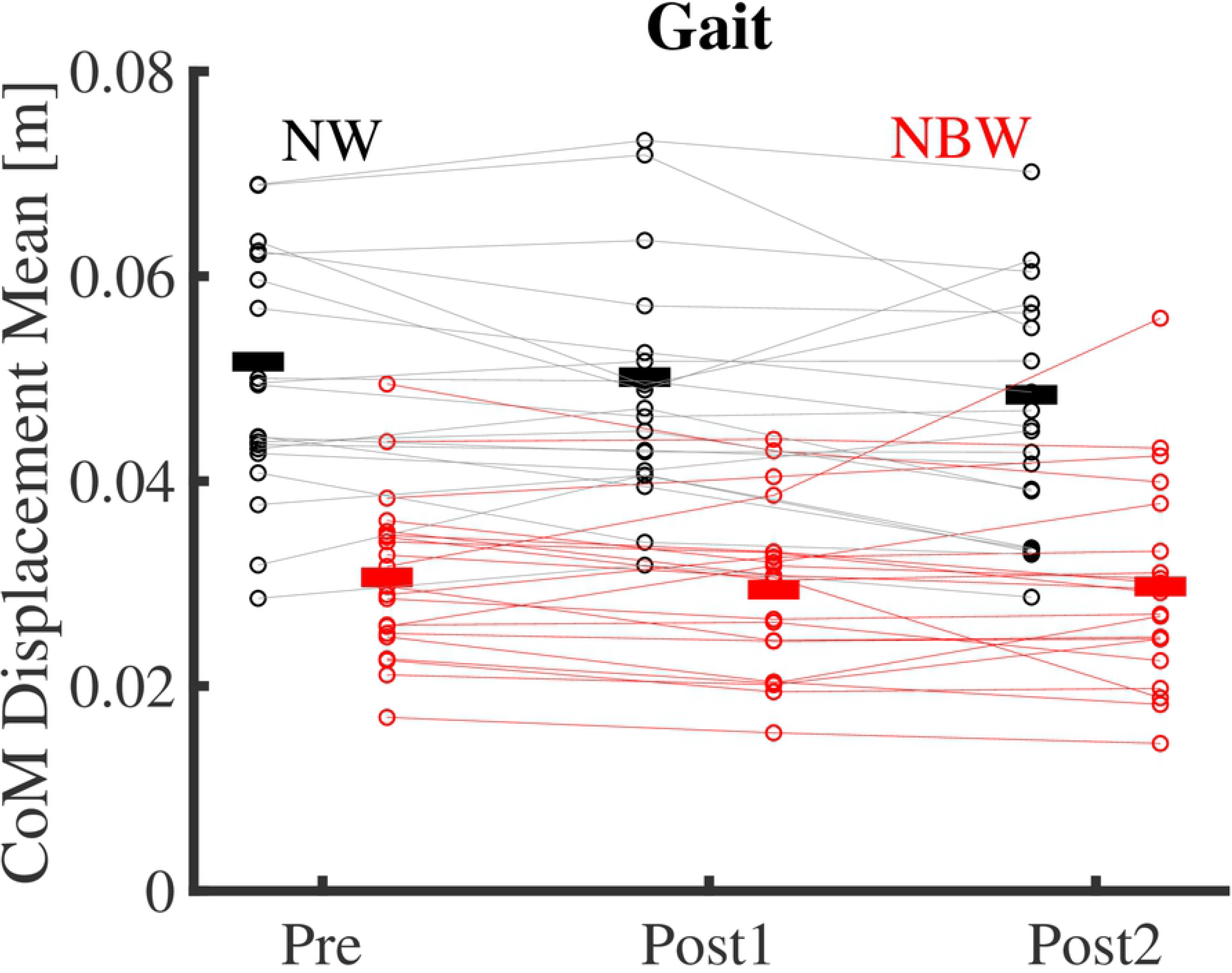
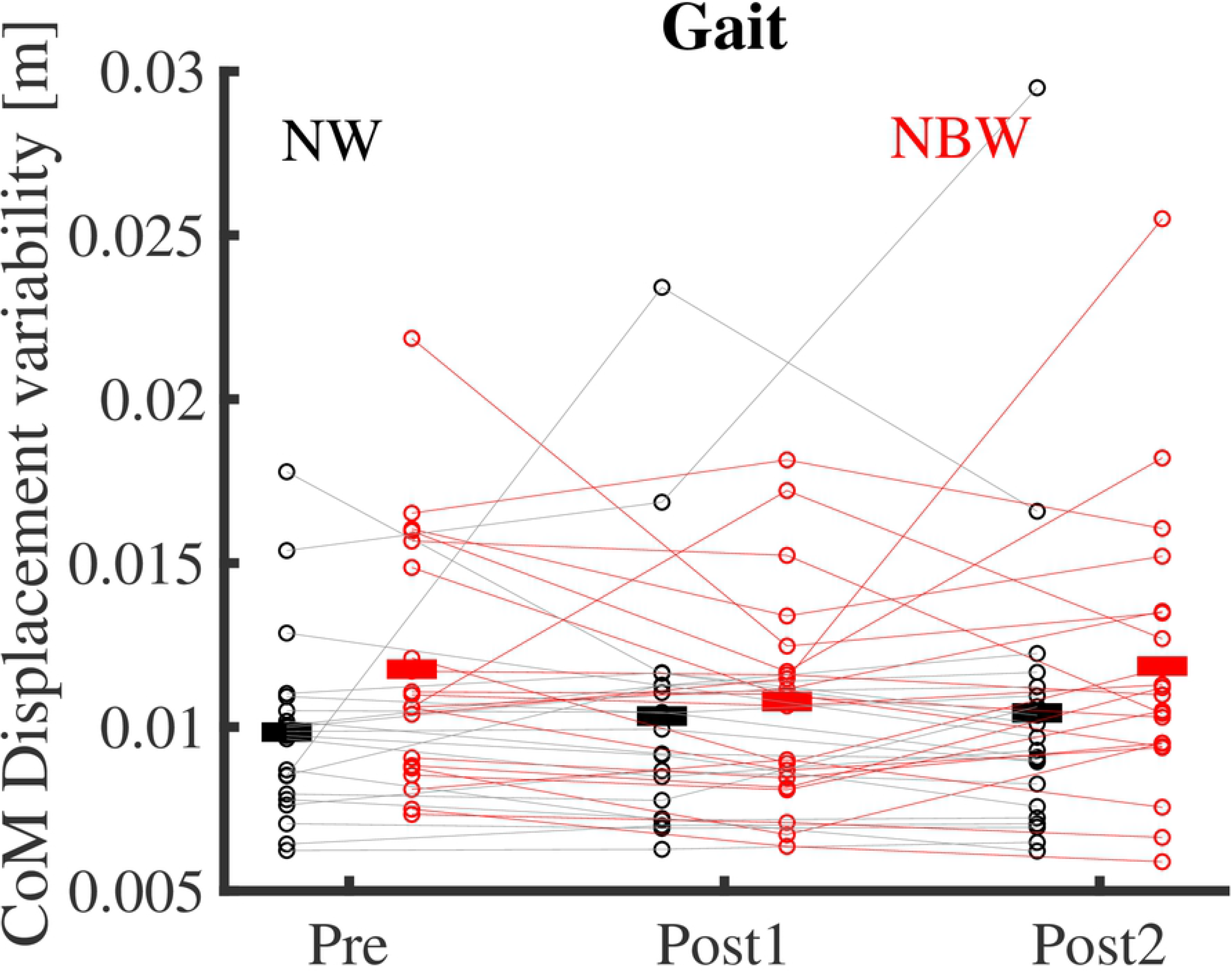
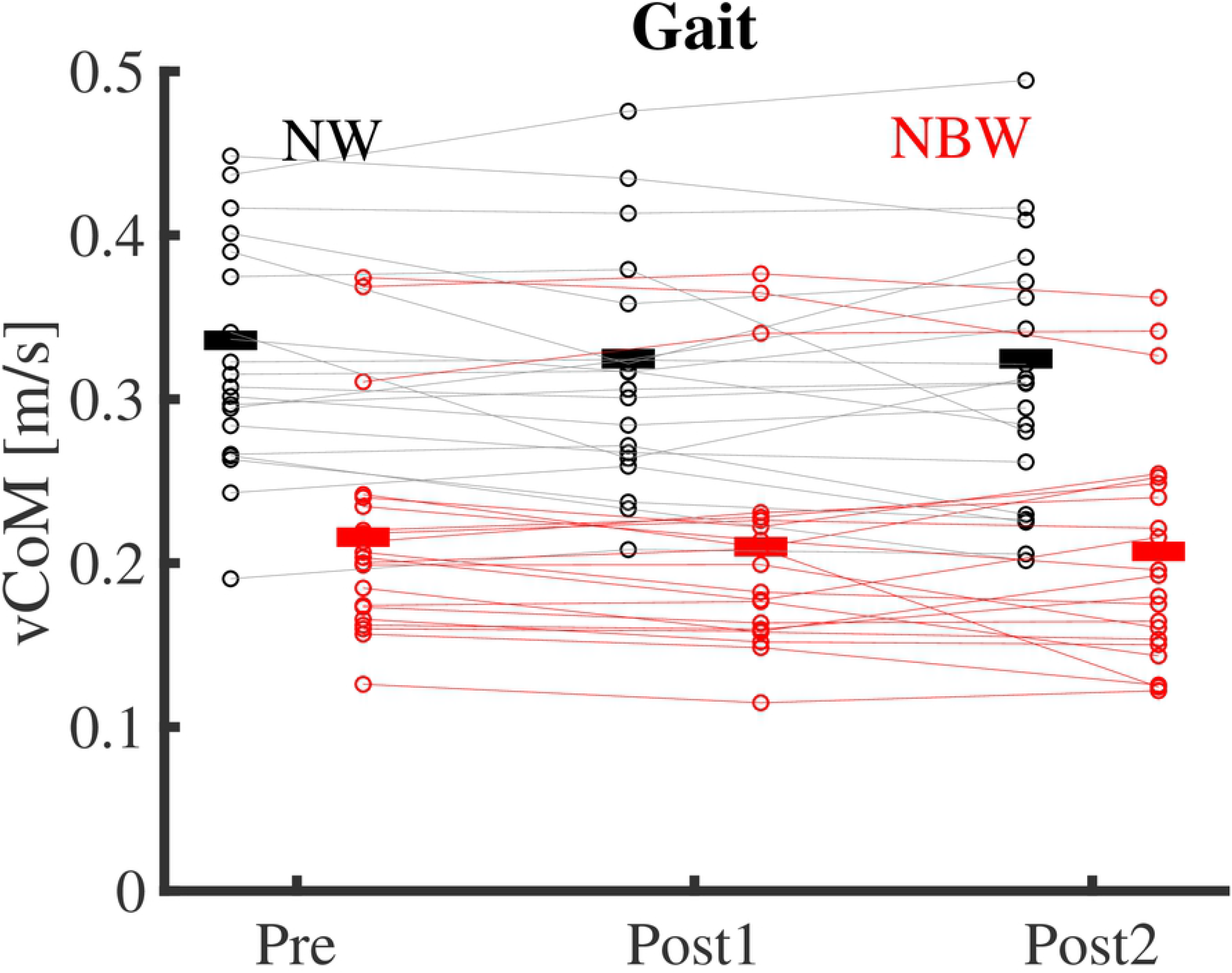
**a)** Mediolateral center of mass displacement and **b)** variability, and **c)** center of mass velocity in narrow-base and normal walking at time-points Pre, Post1 and Post2. Thin lines represent individual subject data. Thick horizontal lines indicate means over subjects. Black; normal walking, red; narrow-base walking.

Training did not significantly affect LDE (F_2,40=_ 0.205, p = 0.814), but condition did, with lower values in narrow-base compared to normal walking (F_1,20=_ 26.223, p < 0.001, Fig 5). No significant interaction of time-point x condition was found (F_1.3,24.699=_ 3.112, p = 0.078).

**Fig 5.**
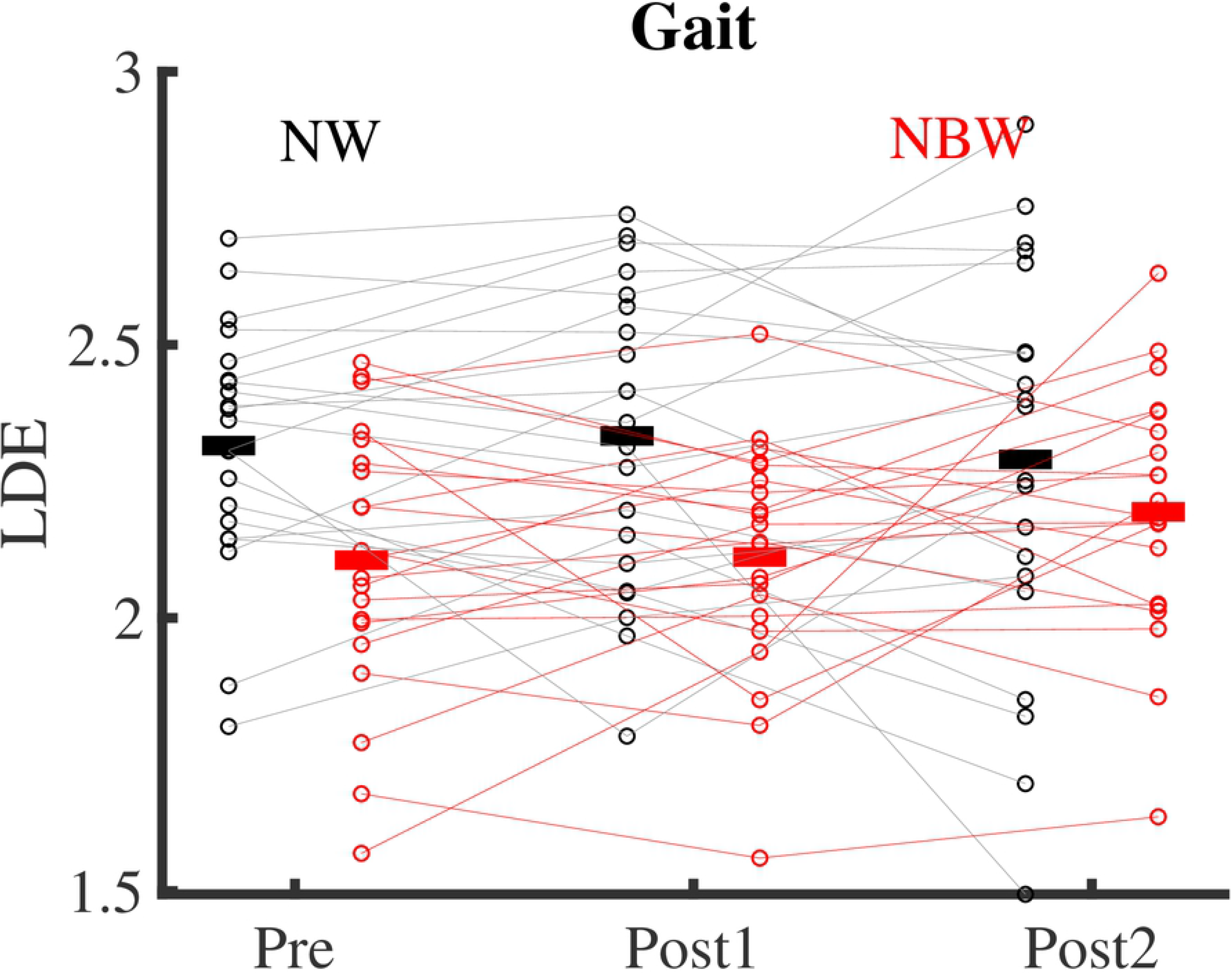
Local divergence exponents in narrow-base and normal walking at time-points Pre, Post1 and Post2. Thin lines represent individual subject data. Thick horizontal lines indicate means over subjects. Black; normal walking, red; narrow-base walking.

### Muscle synergies

Five muscle synergies were extracted with a fixed muscle weighting matrix **H** (Fig 6) and activation profiles per individual per condition and time-point (Fig 7). This accounted for 87±2% of the variance in the EMG data.

**Fig 6.**
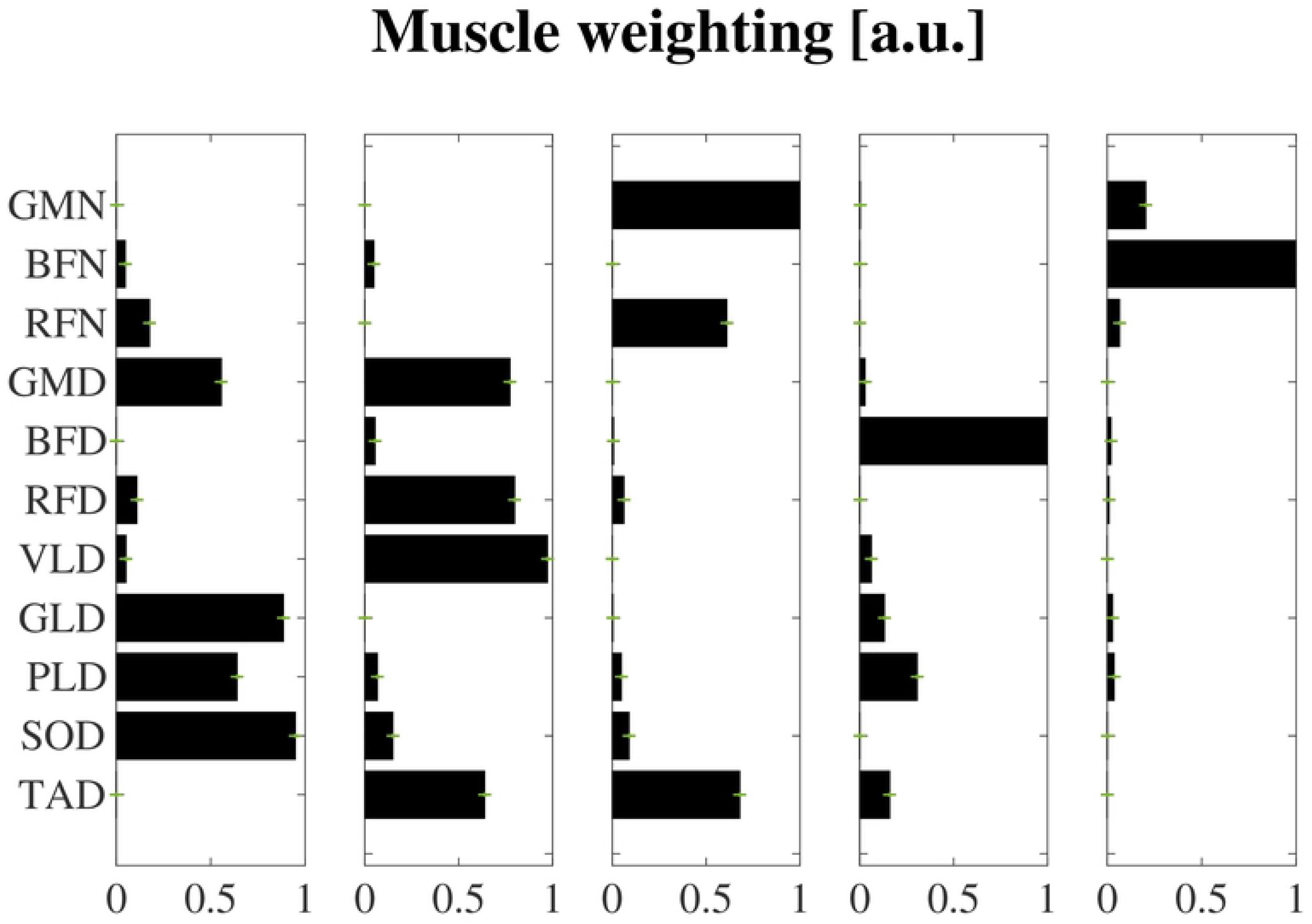
Time- time-invariant muscle weightings of synergies extracted from concatenated data, over all individuals, conditions and time-points. Muscles monitored unilaterally on the dominant side (D): tibialis anterior (TA), vastus lateralis (VL), lateral gastrocnemius (GLD, soleus (SO), peroneus longus (PLD), and muscle collected on the dominant (D) and non-dominant side (N): rectus femoris (RFD, RFN), biceps femoris (BFD, BFN) and gluteus medius (GMD, GMN) muscles.

**Fig7.**
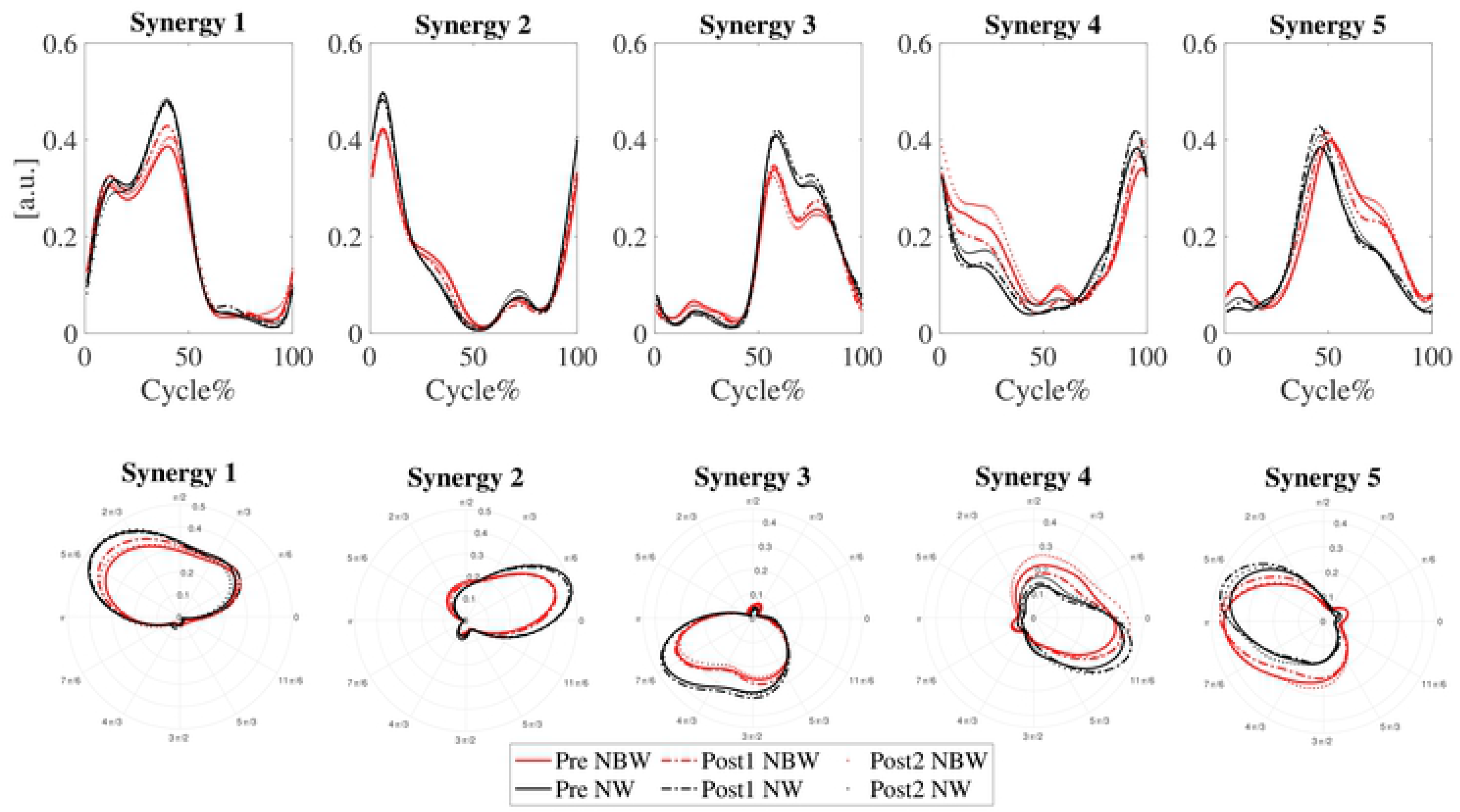
Activation profiles of the extracted synergies as time series and in polar coordinates in narrow-base and normal walking at time-points Pre (solid), Post1 (dash-dot) and Post2 (dotted). The x-axis in the Cartesian coordinates represents one gait cycle. One gait cycle in polar coordinate is [0 2π]. Black; normal walking, red; narrow-base walking.

Based on muscle weightings and activation profiles, the first synergy appeared to be functionally relevant in the stance phase of the dominant leg, with major involvement of soleus and gastrocnemius lateralis. The second synergy appeared to be related to the weight acceptance phase of the dominant leg, where the quadriceps (vastus lateralis, rectus femoris) muscles were mostly engaged. The third synergy resembled partial mirror images of synergies 1 and 2 for the non-dominant leg, but differed due to the fact that only a subset of muscles was measured. It was mainly active in the non-dominant leg’s stance phase, with major involvement of gluteus medius and rectus femoris. It lacks muscle activation related to push-off (represented in synergy 1), because lower leg muscles were not measured and represented thigh muscle activity related to weight acceptance (represented in synergy 2). The fourth synergy appeared to anticipate dominant leg heel-strike with engagement mostly of the dominant leg’s biceps femoris. Finally, the fifth synergy appeared to be the mirror image of the fourth synergy, with pronounced engagement of the biceps femoris of the non-dominant leg.

### FWHM

None of the FWHMs were significantly affected by training. FWHMs were found to be smaller in narrow-base compared to normal walking in the synergies associated with weight acceptance of the dominant leg and the stance phase of the non-dominant leg (synergies 2 & 3; (F_1,20=_ 92.86, p < 0.001; F_1,20=_ 17.06, p < 0.001, respectively, Fig 8). In contrast, FWHM of synergies associated with heel strike appeared to be greater in narrow-base compared to normal walking, but only significantly so for the non-dominant leg (synergies 4 & 5, F_1,20=_ 2.198, p = 0.153; F_1,20=_ 8.603, p = 0.008 respectively, Fig 8). In none of the synergies, FWHM was significantly affected by the interaction of time-point x condition (P > 0.05).

**Fig 8.**
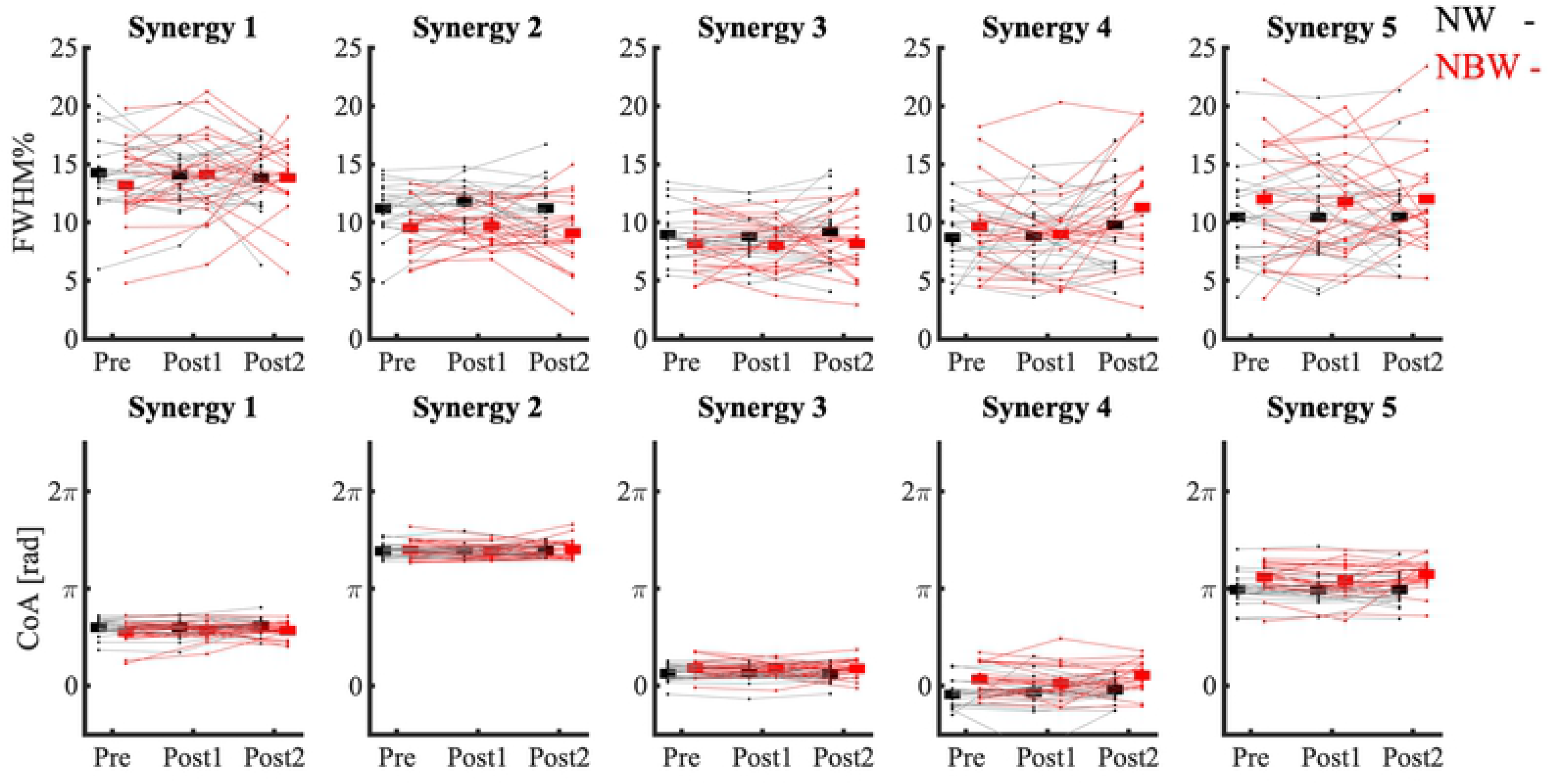
FWHM and CoA of five synergies, in narrow-base and normal walking at time-points Pre, Post1 and Post2. Thin lines represent individual subject data. Thick horizontal lines indicate means over subjects. Black; normal walking, red; narrow-base walking.

### CoA

None of the CoAs were significantly affected by training (p > 0.05). CoA of synergy 1, associated with dominant leg stance, occurred significantly earlier in narrow-base compared to normal walking (F_1,20=_ 6.005, p = 0.015, Fig 8). CoAs of synergy 3 associated with non-dominant stance leg and synergies 4 and 5, associated with heel strike, were delayed in narrow-base compared to normal walking (F_1,20=_ 9.832, p = 0.002; F_1,20=_ 22.109, p < 0.001; F_1,20=_ 18.308, p < 0.001, respectively, Fig 8).

## Discussion

We investigated the transfer of the effects of standing balance training to gait control, by studying gait adaptations to narrow-base walking. We previously reported improvements in robustness and performance of standing balance after short- and long-term standing balance training [24], but here we found no improvements due to training in foot placement error and CoM kinematics during normal or narrow-base walking. Participants adapted their CoM kinematics to foot placement constraints, despite not managing to step consistently within the virtual beam. These adaptations to narrow-base walking did not show an interaction with training. Furthermore, participants adapted to narrow-base walking by modifying activation profiles of their synergies. Standing balance training did not affect these activation profiles, nor their adaptation to narrow-base walking.

In line with literature [8], our participants appeared to control CoM movements more tightly during narrow-base walking than during normal walking, as reflected in a lower CoM displacement and velocity. However, again in line with literature [8], variability of CoM displacement was larger in narrow-base walking. This larger variability might reflect on-line corrections of the CoM trajectory to match it to the constrained foot placement. Confronted with a narrower base, older adults reduced mediolateral CoM displacement and velocity more than young adults [8]. This stronger response might be caused by more cautious behavior, and apparently our balance training did not alter it. Possibly, gait training has more potential to affect balance confidence in gait [40].

Five synergies described leg muscle activity across narrow-base and normal walking, together accounting for 87% of the variation in muscle activity. In spite of differences in muscles measured, participant age and walking conditions between studies, (the number of) these synergies resemble results reported in previous literature [10,41–45]. In our analysis, we kept the muscle weighting in these synergies, constant between conditions and time-points. Participants adapted the activation profiles of these synergies to the gait condition, but no effects of training were observed.

The FWHM of the activation profiles were different between conditions but were not affected by training. An increase of FWHM has been suggested to increase the robustness of gait [20], but in narrow-base walking our participants only increased the FWHM of the activation profile associated with non-dominant leg heel strike (synergy 5), although a similar tendency could be observed for the dominant leg (synergy 4). These adaptations of the activation profiles may reflect increased activity to enhance control over foot placement or to enhance robustness of the new stance leg in preparation for weight transfer. In contrast, participants shortened the FWHM of the activation profiles associated with the stance phase of the non-dominant leg and weight acceptance of the dominant leg. These synergies share muscle activation related to weight acceptance and the change in the activation profiles is mainly visible in a slower build-up of muscle activity (Fig 7). This may reflect a slower weight acceptance by the new support leg, possibly related to the lower activation peak during push-off observable in synergy 1.

The CoA of the activation profiles was different between conditions but was not affected by training. Narrowing step width led to an earlier CoA of the activation profile associated with dominant leg stance (synergy 1) and delayed CoAs of the activation profile associated with dominant and non-dominant leg heel strikes (synergies 4 and 5). Earlier CoA in the dominant leg stance phase appears to be a consequence of the reduction in activation during the second peak of the activation profile (Fig 7). This reduction in activation would reflect a decrease in muscle activity related to push-off and possibly reflects a more cautious gait. The earlier CoA of the activation profile associated with heel strike reflects a more sustained activation following a slower build-up (Fig 7). Again, this may be related to a more cautious walking but also to active control over CoM movement during the stance phase. The latter is supported by the fact that muscles that would contribute to mediolateral control, specifically tibialis anterior, peroneus longus and gluteus medius are part of these synergies. To check that changes in CoA and FWHM of the activation profiles were not due to changes in duration of gait phases, we assessed single support and double support times as percentages of the stride times and no effects of condition were found.

We studied effects of a balance training program of only 3-weeks. For transfer of acquired skills to a new task, it may be necessary that a high skill level is achieved and possibly more than 3 weeks are needed. Improved gait parameters were reported after 12 weeks of balance training [5]. Therefore, a longer duration of training might have led to changes in mediolateral gait stability.

In conclusion, older adults adapted mediolateral CoM kinematics during gait to narrow-base walking and this was associated with changes in synergies governing the activation of leg muscles. However, we found no evidence of a change in control of mediolateral gait stability, nor of these adaptations as a result of balance training.

## Acknowledgment

The research team would like to thank the individuals who participated in the experiment.

## References

1. Rubenstein LZ, Josephson KR, Robbins AS. Falls in the nursing home. Ann Intern Med. 1994;121: 442–451. doi:10.7326/0003-4819-121-6-199409150-00009

2. Berg WP, Alessio HM, Mills EM, Tong C. Circumstances and consequences of falls in independent community-dwelling older adults. Age Ageing. 1997;26: 261–268. doi:10.1093/ageing/26.4.261

3. Giboin L-S, Gruber M, Kramer A. Task-specificity of balance training. Hum Mov Sci. 2015;44: 22–31.

4. Granacher U, Muehlbauer T, Bridenbaugh S, Bleiker E, Wehrle A, Kressig RW. Balance training and multi-task performance in seniors. Int J Sports Med. 2010;31: 353–358. doi:10.1055/s-0030-1248322

5. Martínez-Amat, Antonio; Hita-Contreras, Fidel; Lomas-Vega, Rafael; Caballero-Martínez, Isabel; Alvarez, Pablo J.; Martínez-López E. Effects of 12-week proprioception training program on postural stability, gait, and balance in older adults: A controlled clinical trial. J Strength Cond. 2013;27: 2180–2188.

6. Toebes MJP, Hoozemans MJM, Furrer R, Dekker J, Van Dieën JH. Local dynamic stability and variability of gait are associated with fall history in elderly subjects. Gait Posture. 2012;36: 527–531. doi:10.1016/j.gaitpost.2012.05.016

7. Hof AL, Gazendam MGJ, Sinke WE. The condition for dynamic stability. J Biomech. 2005;38: 1–8. doi:10.1016/j.jbiomech.2004.03.025

8. Arvin M, Mazaheri M, Hoozemans MJM, Pijnappels M, Burger BJ, Verschueren SMP, et al. Effects of narrow base gait on mediolateral balance control in young and older adults. J Biomech. 2016;49: 1264–1267. doi:10.1016/j.jbiomech.2016.03.011

9. Kluft N, Bruijn SM, Luu MJ, Dieën JH va., Carpenter MG, Pijnappels M. The influence of postural threat on strategy selection in a stepping-down paradigm. Sci Rep. 2020;10: 1–9. doi:10.1038/s41598-020-66352-8

10. Bizzi E, Cheung VCK, d’Avella A, Saltiel P, Tresch M. Combining modules for movement. Brain Res Rev. 2008;57: 125–133. doi:10.1016/j.brainresrev.2007.08.004

11. D’Avella A, Saltiel P, Bizzi E. Combinations of muscle synergies in the construction of a natural motor behavior. Nat Neurosci. 2003;6: 300–308. doi:10.1038/nn1010

12. Ivanenko YP, Cappellini G, Poppele RE, Lacquaniti F. Spatiotemporal organization of α‐motoneuron activity in the human spinal cord during different gaits and gait transitions. Eur J Neurosci. 2008;27: 3351–3368.

13. Janshen L, Santuz A, Ekizos A, Arampatzis A. Fuzziness of muscle synergies in patients with multiple sclerosis indicates increased robustness of motor control during walking. Sci Rep. 2020;10: 1–14. doi:10.1038/s41598-020-63788-w

14. Chvatal SA, Ting LH. Common muscle synergies for balance and walking. Front Comput Neurosci. 2013;7: 1–14. doi:10.3389/fncom.2013.00048

15. Baggen RJ, Dieën JH va., Roie E Van, Verschueren SM, Giarmatzis G, Delecluse C, et al. Age-related differences in muscle synergy organization during step ascent at different heights and directions. Appl Sci. 2020;10. doi:10.3390/app10061987

16. da Silva Costa AA, Moraes R, Hortobágyi T, Sawers A. Older adults reduce the complexity and efficiency of neuromuscular control to preserve walking balance. Exp Gerontol. 2020; 111050.

17. Allen JL, Franz JR, Hill C, Carolina N. Control of Movement The motor repertoire of older adult fallers may constrain their response to balance perturbations. 2020; 2368–2378. doi:10.1152/jn.00302.2018

18. Monaco V, Ghionzoli A, Micera S. Age-Related Modifications of Muscle Synergies and Spinal Cord Activity During Locomotion. J Neurophysiol. 2010;104: 2092–2102. doi:10.1152/jn.00525.2009

19. d’Avella A. Modularity for motor control and motor learning. Advances in Experimental Medicine and Biology. Springer, Cham; 2016. pp. 3–19. doi:10.1007/978-3-319-47313-0_1

20. Santuz A, Ekizos A, Eckardt N, Kibele A, Arampatzis A. Challenging human locomotion: Stability and modular organisation in unsteady conditions. Sci Rep. 2018;8: 1–13. doi:10.1038/s41598-018-21018-4

21. Martino G, Ivanenko YP, Avella A, Serrao M, Ranavolo A, Draicchio F, et al. Neuromuscular adjustments of gait associated with unstable conditions. 2020; 2867–2882. doi:10.1152/jn.00029.2015

22. Wang Y, Watanabe K, Asaka T. Effect of dance on multi-muscle synergies in older adults: a cross-sectional study. BMC Geriatr. 2019;19: 340. doi:10.1186/s12877-019-1365-y

23. Sawers A, Allen JL, Ting LH. Long-term training modifies the modular structure and organization of walking balance control. J Neurophysiol. 2015;114: 3359–3373. doi:10.1152/jn.00758.2015

24. Alizadehsaravi L, Koster R, Muijres W, Maas H, Bruijn SM, van Dieen JH. The underlying mechanisms of improved balance after short- and long-term training in older adults. bioRxiv. 2020; 2020.10.01.322313. doi:10.1101/2020.10.01.322313

25. Kluft N, van Dieën JH, Pijnappels M. The degree of misjudgment between perceived and actual gait ability in older adults. Gait Posture. 2017;51: 275–280. doi:10.1016/j.gaitpost.2016.10.019

26. Hilliard MJ, Martinez KM, Janssen I, Edwards B, Mille ML, Zhang Y, et al. Lateral Balance Factors Predict Future Falls in Community-Living Older Adults. Arch Phys Med Rehabil. 2008;89: 1708–1713. doi:10.1016/j.apmr.2008.01.023

27. Maki BE, Holliday PJ, Topper AK. A prospective study of postural balance and risk of falling in an ambulatory and independent elderly population. Journals Gerontol. 1994;49: M72–M84. Available: http://ovidsp.ovid.com/ovidweb.cgi?T=JS&PAGE=reference&D=emed3&NEWS=N&AN=1994105124

28. Sherrington C, Tiedemann A, Fairhall N, Close JCT, Lord SR. Exercise to prevent falls in older adults: an updated meta-analysis and best practice recommendations. N S W Public Health Bull. 2011;22: 78–83. doi:10.1071/nb10056

29. Cappozzo A, Catani F, Della Croce U, Leardini A. Position and orientation in space of bones during movement: anatomical frame definition and determination. Clin Biomech. 1995;10: 171–178.

30. Hermens HJ, Freriks B, Disselhorst-Klug C, Rau G. Development of recommendations for SEMG sensors and sensor placement procedures. J Electromyogr Kinesiol. 2000;10: 361–374. doi:10.1016/S1050-6411(00)00027-4

31. Roerdink M, Coolen BH, Clairbois BHE, Lamoth CJC, Beek PJ. Online gait event detection using a large force platform embedded in a treadmill. J Biomech. 2008;41: 2628–2632. doi:10.1016/j.jbiomech.2008.06.023

32. Hurt CP, Rosenblatt N, Crenshaw JR, Grabiner MD. Variation in trunk kinematics influences variation in step width during treadmill walking by older and younger adults. Gait Posture. 2010;31: 461–464. doi:10.1016/j.gaitpost.2010.02.001

33. Bruijn SM, Van Dieën JH, Daffertshofer A. Beta activity in the premotor cortex is increased during stabilized as compared to normal walking. Front Hum Neurosci. 2015;9: 1–13. doi:10.3389/fnhum.2015.00593

34. Rosenstein MT, Collins JJ, De Luca CJ. A practical method for calculating largest Lyapunov exponents from small data sets. Phys D Nonlinear Phenom. 1993;65: 117–134. doi:https://doi.org/10.1016/0167-2789(93)90009-P

35. Bruijn S. SjoerdBruijn/LocalDynamicStability: First release. 2017 [cited 2 Sep 2020]. doi:10.5281/ZENODO.573285

36. Oliveira ASC, Gizzi L, Kersting UG, Farina D. Modular organization of balance control following perturbations during walking. J Neurophysiol. 2012;108: 1895–1906. doi:10.1152/jn.00217.2012

37. Cappellini G, Ivanenko YP, Martino G, MacLellan MJ, Sacco A, Morelli D, et al. Immature spinal locomotor output in children with cerebral palsy. Front Physiol. 2016;7: 1–21. doi:10.3389/fphys.2016.00478

38. Santuz A, Ekizos A, Janshen L, Baltzopoulos V, Arampatzis A. The influence of footwear on the modular organization of running. Front Physiol. 2017;8. doi:10.3389/fphys.2017.00958

39. Harrison D, Kanji GK. The development of analysis of variance for circular data. J Appl Stat. 1988;15: 197–223. doi:10.1080/02664768800000026

40. Park S-K, Kim S-J, Yong Yoon T, Lee S. Effects of circular gait training on balance, balance confidence in patients with stroke : a pilot study. J Phys Ther Sci. 2018;30: 685–688.

41. Ivanenko YP, Poppele RE, Lacquaniti F. Five basic muscle activation patterns account for muscle activity during human locomotion. 2004;1: 267–282. doi:10.1113/jphysiol.2003.057174

42. Ghionzoli A, Micera S. Age-Related Modifications of Muscle Synergies and Spinal Cord Activity During Locomotion. 2020; 2092–2102. doi:10.1152/jn.00525.2009.

43. Clark DJ, Va MR. Motor modules of human locomotion : influence of EMG averaging, concatenation, and number of step cycles. 2014. doi:10.3389/fnhum.2014.00335

44. Cappellini G, Ivanenko YP, Poppele RE, Lacquaniti F. Motor patterns in human walking and running. J Neurophysiol. 2006;95: 3426–3437. doi:10.1152/jn.00081.2006

45. van den Hoorn W, Hodges PW, van Dieen JH, Hug F. Effect of acute noxious stimulation to the leg or back on muscle synergies during walking. J Neurophysiol. 2015;113: 244–254. doi:10.1152/jn.00557.2014

